# Walking (and talking) the plank: Dual-task performance costs in a virtual balance-threatening environment

**DOI:** 10.1101/2023.10.29.564612

**Authors:** Tiphanie E. Raffegeau, Sarah A. Brinkerhoff, Mindie Clark, Ashlee D. McBride, A. Mark Williams, Peter C. Fino, Bradley Fawver

**Author notes:** **Disclaimer:** Material has been reviewed by the Walter Reed Army Institute of Research. There is no objection to its presentation and/or publication. The opinions or assertions contained herein are the private views of the author, and are not to be construed as official, or as reflecting true views of the Department of the Army or the Department of Defense. The investigators have adhered to the policies for protection of human subjects as prescribed in AR 70–25. **Corresponding Author:** Tiphanie E Raffegeau, PhD Assistant Professor George Mason University 10890 George Mason Circle Katherine Johnson Hall 201G, MSN 4E5 Manassas, VA 20110.

## Abstract

We evaluated the effects of engaging in extemporaneous speech while walking in virtual environments meant to elicit low or high levels of mobility-related anxiety. We expected that mobility-related anxiety imposed by a simulated balance threat (i.e. virtual high elevation) would impair walking behavior and lead to greater dual-task costs. Altogether, 15 adults (age = 25.6 ± 4.7 yrs, 7 women) walked at their self-selected speed within low (ground) and high elevation (15 meters) VR settings while speaking extemporaneously (dual-task) or not speaking (single-task). Likert-scale ratings of cognitive and somatic anxiety, confidence, and mental effort were evaluated after experiencing each condition, and gait speed, step length, and step width, and the variability of each, was calculated for each trial using the position of trackers attached to participants’ ankles. Silent speech pauses (>150ms) were determined from audio recordings to infer the cognitive costs of extemporaneous speech planning at low and high virtual elevation. The presence of a balance threat and the inclusion of a concurrent speech task both perturbed gait kinematics, but only the virtual height illusion increased anxiety and mental effort while decreasing confidence. Extemporaneous speech pauses were longer on average when walking, but no effects of virtual elevation were reported. Trends toward interaction effects arose in self-reported responses, participants reported more comfort walking at virtual heights if they engaged in extemporaneous speech. Walking at virtual elevation and walking while talking have independent and significant effects on gait; both effects were robust and did not support an interaction when combined (i.e., walking and talking at virtual heights). Rather than additive cognitive-motor demands, the nature of extemporaneous speech may have distracted participants from the detrimental effects of walking in anxiety-inducing settings.

## 1. Introduction

Mobility in daily life often requires managing concurrent cognitive and motor demands under conditions that can threaten balance, such as talking to a friend while navigating a busy crosswalk. The influence of cognitive and motor demands on mobility behavior has become a particular focus within psychology and the movement sciences that describe how interference between cognitive and motor processes influence mobility behavior under various environmental and contextual demands.^1,2^ Researchers have specifically sought to understand how individuals manage simultaneous perceptual-cognitive demands during locomotor tasks in young adults,^3,4^ older adults,^5–7^ in people with deficits in cognitive-motor function,^8–10^ and patients with movement disorders like Parkinson’s disease,^11–13^ focusing on factors influencing fall-risk. However, empirical studies tend to ignore individual differences in environmental risk-assessment and the reciprocal influence of mobility-related anxiety on cognitive-motor behaviors.^2,14^

Theoretical frameworks that address the hierarchical nature of dual-tasking during gait suggest that healthy people use a ‘posture-first’ strategy by focusing primarily on their gait performance in hazardous situations where balance is at risk.^2^ The integrated prioritization model further implies that individual differences in relative motor (i.e., ‘postural reserve’) and cognitive capabilities (i.e. ‘cognitive reserve’) dictate the allocation of attention during gait.^2^ Attentional Control Theory (ACT) provides an alternative framework to understand how performance-related state anxiety influences perceptual-cognitive control.^14,15^ ACT specifies that anxiety disrupts attentional control through increased bottom-up ‘distraction bias’ to external threats in lieu of processing goal-directed (i.e., task-relevant) information.^14,15^ When applied to walking performance, ACT predicts that an individual undergoing acute fall-related anxiety would likely disengage from any additional cognitive demand as perceptual-cognitive resources are already strained by the addition of worrisome thoughts about performance, such as self-preoccupation and concerns about performance evaluation.^14^ Within the context of posture and gait behavior, other related frameworks such as Conscious Processing Theory ^16^ operationalize disruptions in attentional and motor control as ‘reinvestment,’ which entails devoting resources to a task that was previously autonomous. Reinvestment can lead to sub-optimal cognitive and sensory functioning during walking,^17–19^ as well as promote ‘self-focus’ or ‘internal-focus’ ^20^ that can inhibit processing of external stimuli during complex mobility tasks for older adults.^19,21^ Existing theories suggest that cognitive demands (e.g., a dual task) and psychological demands (e.g., brought on by balance threat) draw from the same limited pool of cognitive resources. Competition for shared resources is thought to lead to rigidly controlled motor behavior (e.g., reinvestment in controlled processes, internal focus), or distraction from motor skill execution (e.g., through task-irrelevant thoughts, increased sensitivity to external stimuli). ^17–21^

A well-practiced cognitive task that does not directly compete with sensory integration may elicit different effects on cognitive-motor control in stressful environments, but most existing research has typically relied on observations from tasks that might be influenced by age-related declines in sensory interference, such as an auditory reaction time task ^22^ or a visuospatial distractor (i.e. clock-monitoring).^23^ In healthy adults, published reports have highlighted conflicting sensorimotor goals between auditory ^3,24^ or visually demanding ^25^ cognitive tasks and visual integration for gait. Many researchers attempting to avoid sensory interference have used the serial subtraction task (i.e., subtract from 100 by 3 or 7),^26,27^ which challenges cognitive processes but is subject to biases associated with socio-economic background or education levels ^28^ and can impose a rhythmicity on gait that could influence walking performance.^29,30^ Moreover, contrived cognitive tasks have a ‘purity’ problem, in which a targeted cognitive process engages broader network-wide processes.^31^ Therefore, such contrived cognitive tasks may serve as distractions that bias attentional control more than well-practiced cognitive demands, particularly in anxiety-inducing settings. Finally, due to learning effects,^32^ there is concern over whether results derived from previous studies that average performance across multiple trials represent realistic cognitive-motor behavior that does not involve repeated practice. Alternatively, the social consequences of extemporaneous speech production can engage healthy adults with speech production demands such that talking behavior is prioritized over gait performance.^4^ We have previously observed that only when the demands of a motor task become too difficult (i.e. avoiding an obstacle) will healthy young adults demonstrate a trade-off between concurrent extemporaneous speech and complex locomotion; allowing costs to speech production in favor of dedicating resources to motor performance when the locomotor task is demanding.^4^ It is feasible that walking while talking in contexts that elicit mobility-related anxiety would result in different resource allocation patterns than those previously observed using laboratory-based tasks. By examining the single and dual-task costs associated with well-practiced cognitive demands under conditions of low and high perceived threats to mobility, we may be able to distinguish the influence of relevant cognitive demands on attentional biases in anxiety-inducing settings.

In the present study, we build on existing theoretical frameworks to examine how individuals manage cognitive-motor demands in situations that elicit anxiety about gait performance. Specifically, we used a virtual reality (VR) based approach to induce state anxiety (i.e., walking at a simulated high elevation), as individuals walked alone (single-task) and walked while performing a concurrent extemporaneous speech monologue (dual-task). Aligning with evidence from similar studies in the field ^33–37^ and prevailing theoretical frameworks informing cognitive, perceptual, and motor costs during gait,^18,38,39^ we predicted that walking at virtual elevation without a concurrent cognitive task would be associated with more ‘protective’ walking behavior (i.e., slower gait speed and shorter and wider steps to avoid potential balance perturbations) compared to simulated ground level walking. Since extemporaneous speech is cognitively demanding, but does not directly conflict with sensory demands during walking, we suspected that young healthy adults would not demonstrate a ‘tradeoff’ between cognitive and gait performance. We instead predicted that participants would preserve their speech performance within anxiety-inducing settings, even as gait behavior became more conservative. Conversely, if healthy adults allowed both cognitive and motor performance to decline while walking in anxiety-inducing settings, it would suggest that balance threat leads participants to dedicate resources to prevent a potential fall, irrespective of the threat-related relevance of the cognitive task (i.e., anxiety about walking behavior altering speech behavior). Based on ACT,^15,40^ we predicted that at ground level virtual elevation (i.e., without a balance threat), young adults would prioritize the extemporaneous speech task and exhibit compensations in gait behavior as compared to walking without the cognitive dual-task (i.e., slower gait speed, shorter and wider steps). However, at virtual high elevation, we expected performance costs brought on by greater levels of mobility-related anxiety would be reflected in gait and speech outcomes, such that individuals would adopt conservative gait behavior (i.e., slower gait speed, shorter and wider steps) and exhibit interference in speech performance (i.e., more frequent speech pauses of greater duration).

## 2. Methods

### 2.1 Participants

Healthy participants were recruited using a convenience sampling strategy. Individuals were included if their vision was normal or corrected to normal, they had no orthopedic injuries causing discomfort during walking, and English was their primary language. No participant reported experiencing a fall in the previous six months, defined as ‘coming to a lower level unintentionally.’ ^41^ All participants provided informed consent using a protocol approved by the local Institutional Review Board.

### 2.2 Measures

#### 2.2.1 Dispositional Anxiety

Anxiety was assessed using the State-Trait Anxiety Inventory (STAI).^42^ Scores range from 20-80, with higher scores reflecting greater trait (i.e., generally) or state (i.e., day of the study) anxiety.

#### 2.2.2 Cognitive Function

Participants completed two common clinical tests of cognition (i.e., executive functioning) on paper, the Stroop Task and the Trial Making Test, both of which have been shown to be strongly related to cognitive-motor performance and mobility.^43,44^ Our version of the Stroop Task consisted of two parts: (1) Congruent, a test of response time, requiring participants to name as many colored letters (e.g., “XXXXX”) as possible within 45 seconds (measuring visual scanning and response time); and (2) Incongruent, a measure of response inhibition (measuring executive function maintenance and set switching), requiring participants to name as many ink colors of mismatched color-words in 45 seconds.^45,46^ Three participants reported being red-blue colorblind and did not complete the Stroop tests.

#### 2.2.3 Perceptions of Performance

We used validated visual analog scales to capture self-reported perceptions of performance. The Mental Readiness Form-3 (MRF-3)^47^ was administered to assess cognitive (i.e., worry) and somatic (i.e., arousal) components of anxiety experienced during each condition, as well as participants’ level of confidence in their ability to complete the task, using an 11-point Likert-scale. Cognitive anxiety ratings (i.e., worrying thoughts) were prompted with the root statement ‘my thoughts were’, with responses ranging from 1, ‘very calm,’ to 11, ‘very worried.’ Somatic anxiety ratings (i.e., moment to moment changes in physiological arousal) were prompted by the root statement ‘my body feels,’ and ranged from 1, ‘very relaxed,’ to 11, ‘very tense.’ Participants self-rated confidence in their ability to complete the task was evaluated with the root statement ‘I am feeling,’ and response ranging from 1, ‘very confident,’ to 11, ‘not confident at all’. For analysis, we reverse-scored this measure so that higher values would indicate greater levels of confidence in their ability to complete the task. Participants indicated the level of mental effort required to complete the task in each condition using the Rating Scale of Mental Effort (RSME),^48^ which is a 0 to 150 scale that ranges from ‘0: absolutely no effort’ to ‘150: extreme effort.’

#### 2.2.4 Gait Kinematics

Step length, step width, and gait speed, and the variability (standard deviation) of each were calculated using a custom MATLAB script (version R2022b, Natick, MA) by using the linear position from the HMD and the two ankle tracking accessories placed on the lateral aspect of participants’ ankles were recorded for each trial (see Data Processing).

#### 2.2.5 Speech Performance

The participant was fitted with a wireless microphone (Lavalier, model WMX-1) to record speech. The number and mean duration of pauses during the extemporaneous speech task (defined as a speech pause > 150 milliseconds) were identified by a trained research assistant using open access software (PRAAT, v 6.2.14).

### 2.3 Instrumentation and Task

#### 2.3.1 Virtual Gait Task

Participants wore the HTC Vive (version 2.0, Bellevue, WA) immersive head mounted display (HMD) displaying a 0.4 x 5.2m virtual walkway matched to a real path. The global 3D coordinates for each corner of the walkway in the virtual space were determined by capturing the position of each corner of the physical walkway using a hand controller, matching the dimensions and coordinates of the real-world walkway to the virtual simulation. In accordance with International Society of Biomechanics Standards, the center and starting point of the walkway was defined as the origin, x was the sagittal path of progression, z was the mediolateral axis, and y was the vertical axis.^49^ Participants wore trackers around their ankles (HTC Vive version 2.0, Bellevue, WA) to provide ongoing visual feedback of where their feet were in the virtual space. For two minutes, participants explored the virtual space and were allowed to walk or stand along the walkway in whatever fashion they preferred. After the familiarization period, participants removed the headset and practice trials without VR, on which data were not collected. To collect participants’ perceptions during each trial block, rating scales were presented inside the headset with similar instructions for completion using a hand controller (Figure 1).

**Figure 1.**
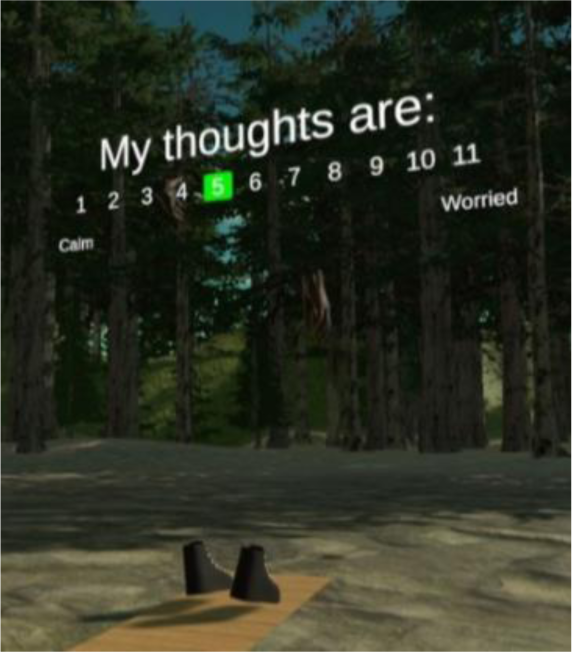
After each condition, participants used a hand controller to select their responses for each self-report item. Visual analog scales were presented in the virtual environment to determine the participants level of somatic and cognitive (shown here) anxiety and confidence (Mental Readiness Form; MRF-3), as well as their level of mental effort devoted to task completion (Rating Scale for Mental Effort; RSME).

#### 2.3.2 Extemporaneous Speech Task

Each participant was offered a list of 26 conversation topics (e.g., first job, favorite TV show, recent trips taken) and asked to choose six of those topics they could talk about for at least one minute. At the beginning of each dual-task trial, participants were randomly assigned one of their selected topics to speak about. Participants first completed the extemporaneous speech task while seated in the laboratory space, representing single-task cognitive performance. The instructions emphasized that ‘what you say doesn’t matter, just that you can keep talking the entire time’.

### 2.4 Procedure

Each data collection session began with the participant completing a series of surveys and cognitive tests to address dispositional differences (Table 1). Virtual walking conditions (i.e., blocks) were pseudorandomized so that the order of single vs. dual task conditions were always counterbalanced across participants. The virtual low elevation environment was always presented first in each single or dual task condition to ensure participants experienced the largest effects of mobility-related anxiety at virtual high elevation (1^st^ Block = Low, 2^nd^ Block = High).^15^ Participants walked continuously at their self-selected pace for one minute, completing between 7-12 passes on the 5.2 meter walkway consisting of 3-5 strides per pass. To transition between low and high virtual elevation, participants were seated in a chair facing forward at the beginning of the walkway with their feet on the path and eyes looking straight ahead as they were lifted to an approximate 15-meters above ground at the rate of a standard elevator (1 m/s). During walking trials, participants were instructed to ‘stay on the path and walk without speaking at their comfortable pace and continue walking until they heard ‘stop’. During dual-task trials, participants were instructed to ‘walk and talk like you were speaking to a friend’ with no additional instructions to prioritize one task over another.

**Table 1.** Participant Characteristics (*N* = 15)

|  | Mean | SD | Range |  |
| --- | --- | --- | --- | --- |
|  |  |  | Lower limit | Upper limit |
| Age (y) | 25.6 | 4.7 | 22.63 | 27.81 |
| Height (cm) | 168.0 | 10.7 | 161.5 | 173.0 |
| Mass (kg) | 68.5 | 12.5 | 59.7 | 77.1 |
| Leg length (mm) | 92.8 | 6.7 | 87.4 | 97.4 |
| Gender (women/men) | 7/8 | -- | -- | -- |
| STAI-T (score) | 31.4 | 4.6 | 31.0 | 38.2 |
| STAI-S (score) | 25.2 | 5.2 | 23.0 | 33.2 |
| Stroop Congruent (score) | 93.6 | 8.1 | 86.5 | 99.5 |
| Stroop Incongruent (score) | 75.6 | 14.5 | 69.5 | 85.5 |
| TMT-A (s) | 17.3 | 5.1 | 12.9 | 21.0 |
| TMT-B (s) | 34.9 | 14.3 | 29.2 | 43.3 |

*Note:* y = years; cm = centimeters; kg = kilograms; mm = millimeters; STAI-T/S = State-Trait Anxiety Inventory; TMT-A/B = Trails Making Test A and B; SD = standard deviation; Stroop scores represent the number of correct responses per 45 seconds; s = seconds.

### 2.5 Data Processing

The HTC Vive collects variable position sampling rates (e.g., one trial frame rate range = 89Hz to 93Hz) depending on the relative speed of motion and the independent sampling rates of the lighthouses and tracker accessories.^16^ Consequently, we resampled the data to 100Hz using the *resample* function with linear interpolation in MATLAB. Errors in position tracking were identified by removing erroneous position data that were recorded below the origin (the walkway) and replacing resultant missing data with spline-filled data points. The spline-filled data were then filtered with a zero-pass fourth order Butterworth filter. Foot contacts were identified as peaks in the vector between the HMD and each foot tracker. 3D position of the feet and time at each identified peak was extracted at foot strike.^50^

Straight steps were isolated by retaining the steps taken within the central 4.4-meter portion of the walkway and removing turning steps from the gait analysis. However, some individuals did not walk the entire length of the walkway, especially at virtual high elevation. Therefore, for each individual, we calculated their maximum distance travelled on the walkway and extracted steps within 0.3 meters from that individual’s maximum distance. Step length was calculated for each foot contact as the absolute distance between the ankle-worn sensors in the anterior posterior direction, and step width was the absolute distance between the ankle-worn sensors in the mediolateral direction for successive steps. Gait speed was calculated as step length divided by the time between two consecutive footfalls. Left and right step values were averaged to represent overall gait performance. Variability was calculated as the standard deviation across steps.

The frequency and duration of silent speech pauses are interpreted as indicators of cognitive costs incurred in each condition.^51,52^ Previous work has demonstrated speech pauses during a extemporaneous speech monologue are sensitive to motor difficulty.^4^ Seated speech is considered the baseline for cognitive performance capacity (single-task). The number and mean duration of pauses during the extemporaneous speech task (silent pause > 150 milliseconds) were identified by a trained research assistant using open access software (PRAAT, v 6.2.14). Based on previous methods^4,51^ research assistants marked the beginning and end of silent pauses using spectrograms and waveforms. A custom MATLAB code determined pause length, the number of pauses, and the total pause time within a trial.

### 2.6 Statistical Analyses

We used the *fitlme* and *anova* (ANOVA; analysis of variance) functions in MATLAB to analyze linear mixed-effect regression models and type III tests for fixed effects, respectively. Separate, fully factorial linear mixed-effect regressions (LMERs) were used to evaluate the effect of Height (low vs. high) and Cognitive Demand (single vs. dual) on gait performance (i.e., gait speed, step length, step width, and their variability). Models included a fully crossed random intercept by participant, participant within Height, and participant within Cognitive Demand, accounting for within-participant variance at each elevation and cognitive function. Height was reference-coded such that low elevation was the reference (Height: low = 0, high = +1). Cognitive Demand was reference-coded such that single-task was the reference (cognitive demand: single-task = 0, dual-task = +1). Therefore, the reported unstandardized *β* (beta) weights and respective confidence intervals [CI] can interpreted as mean differences between factor levels, and interaction effects would represent change due to virtual elevation multiplied by the change from single to dual task. ANOVA *F*-scores represent a standardized relative effect for each model. All model outputs are provided in the Supplementary Appendix.

Self-reported ratings were analyzed using fully factorial LMERs to determine the effect of Height (low = reference vs. high) and Cognitive Demand (single = reference vs. dual-task) on perceived anxiety (somatic and cognitive), confidence, and mental effort. Although LMER is robust to violations of normality, we bootstrapped the values (*N* = 1000), sampling with replacement, to compare more robust confidence intervals in self-report ratings between low and high virtual elevation. Finally, we used mixed model ANOVAs to determine the effect of Task-Elevation (seated single task = reference vs. low DT vs. high DT) on the number of speech pauses, speech pause length, total speech pause duration. We included a random intercept of subject within condition in all models, and for speech pause length we included a random intercept and slope of subject within condition. The significance threshold for all statistical analyses was set at *α* = 0.05.

## 3. Results

### 3.1 Demographics and Self-Report

Out of an initial 18 participants that were evaluated, we were unable to include data from the first three due to a technical difficulty which was subsequently resolved with a slight change in the protocol related to setting up the VR system. All included participant demographics (*N* = 15) are reported in Table 1. No substantial variability was observed among participant characteristics other than anthropometrics, Stroop Incongruent trials (i.e., response inhibition), and the Trails Making Test-B (i.e., executive function, set switching).

Analyses of self-reported ratings following each condition revealed statistically significant main effects of Height on participants perceptions of the cognitive, *F*(1,56) = 64.32, *p* < 0.001, and somatic, *F*(1,56) = 85.03, *p* < 0.001, components of anxiety during walking trials, as well their confidence, *F*(1,56) = 53.81, *p* < 0.001, and mental effort, *F*(1,56) = 35.60, *p* < 0.001, in executing the experimental task (see Figure 2 and Supplementary Appendix Tables 1-4). Decomposition of these main effects indicated that, relative to the range of possible self-report response values (scored 1-11), walking at virtual elevation resulted in participants experiencing an approximate 31% increase in worrying thoughts (*β* = 3.42 [2.53,4.32]), 30% increase in perception of changes in arousal (*β* = 3.28 [2.55, 4.02]), 26% decrease in their confidence (*β* = -2.88 [-3.77,-1.98]), and a 20% increase in their mental effort (*β* = 29.40 [16.07,42.72]). No main effects of Cognitive Demand (i.e., presence of a dual task) were documented for any self-report outcome (all *p*’s > 0.111). Finally, no statistically significant interactions of Height x Cognitive Demand were observed for self-report items; however, the interaction effects for cognitive anxiety, *F*(1,56) = 10.88, *p* = 0.78, and confidence, *F*(1,56) = 18.02, *p* = 0.73, trended towards significance, suggesting that walking while talking somewhat mitigated the increases in worrying thoughts and decreases in self-efficacy experienced when walking at high elevation (*β* = -0.89 [-0.36,-0.27]).

**Figure 2.**
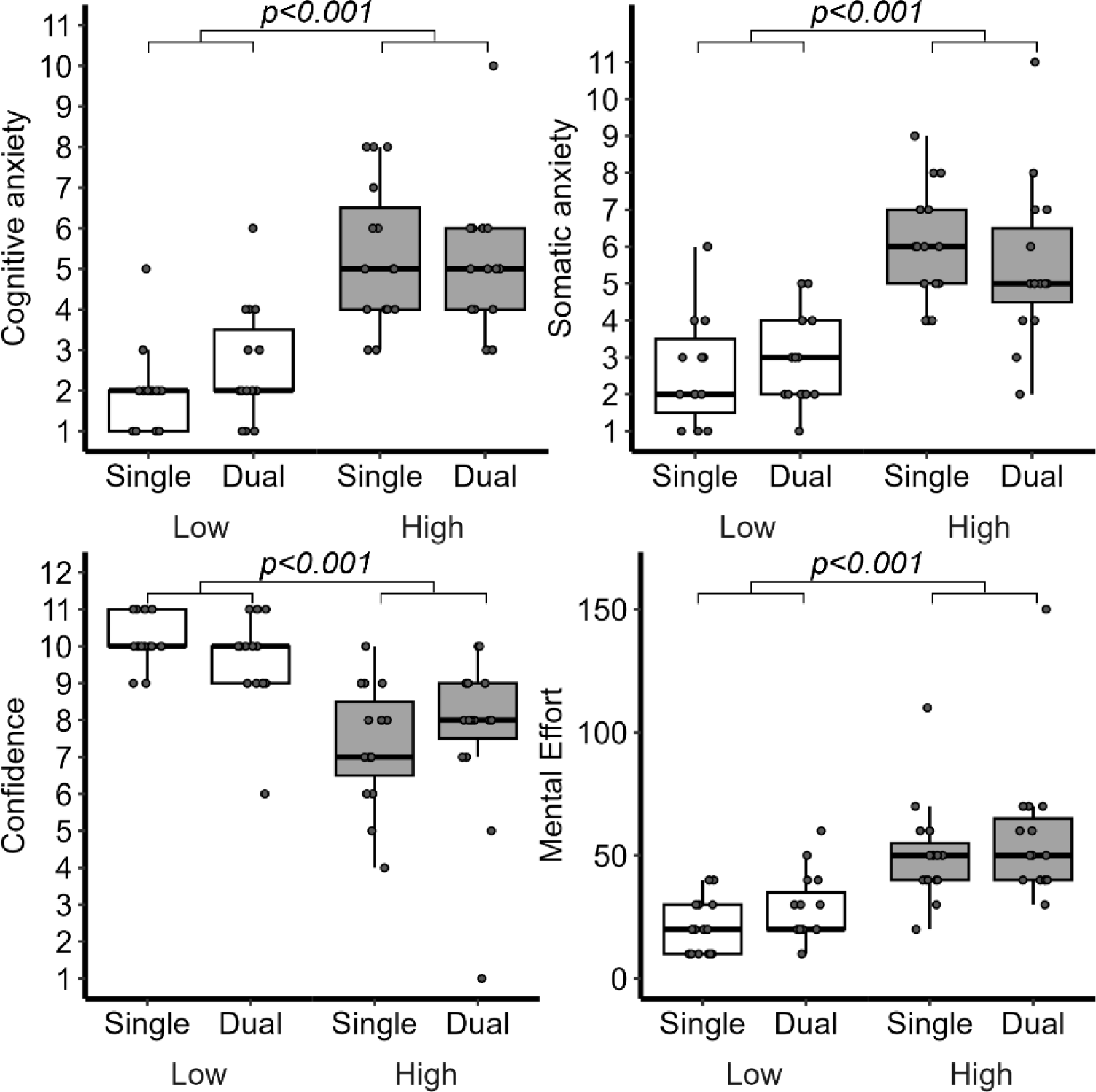
The changes in self-reported Cognitive Anxiety (worry; Top Left), Somatic anxiety (tension; Top Right), Confidence (Bottom Left), and Mental Effort (Bottom Left) across each walking condition.

### 3.2 Gait Performance

There were main effects of Height, *F*(1,52) = 7.15, *p* = 0.010, and Cognitive Demand, *F*(1,52) = 32.11, *p* < 0.001, on gait speed, indicating that participants walked 11% slower during the high elevation condition compared to low elevation (*β* = -0.11 m/s [-0.20,-0.03]), and 14% slower during dual-task compared to single task conditions (*β* = -0.16 m/s [-0.21,-0.10]). No significant interactions were detected for gait speed (*p* = 0.951). Gait speed variability was significantly impacted by the presence of a dual task, *F*(1,52) = 18.21, *p* < 0.001, indicating participants exhibited more consistent walking speed across trials while engaging in extemporaneous speech (*β* = -0.05 m/s [-0.07,-0.03]), but no main effect of Height or interaction was revealed (*p*’s > 0.696; see Figure 3 and Supplementary Tables 5-6).

**Figure 3.**
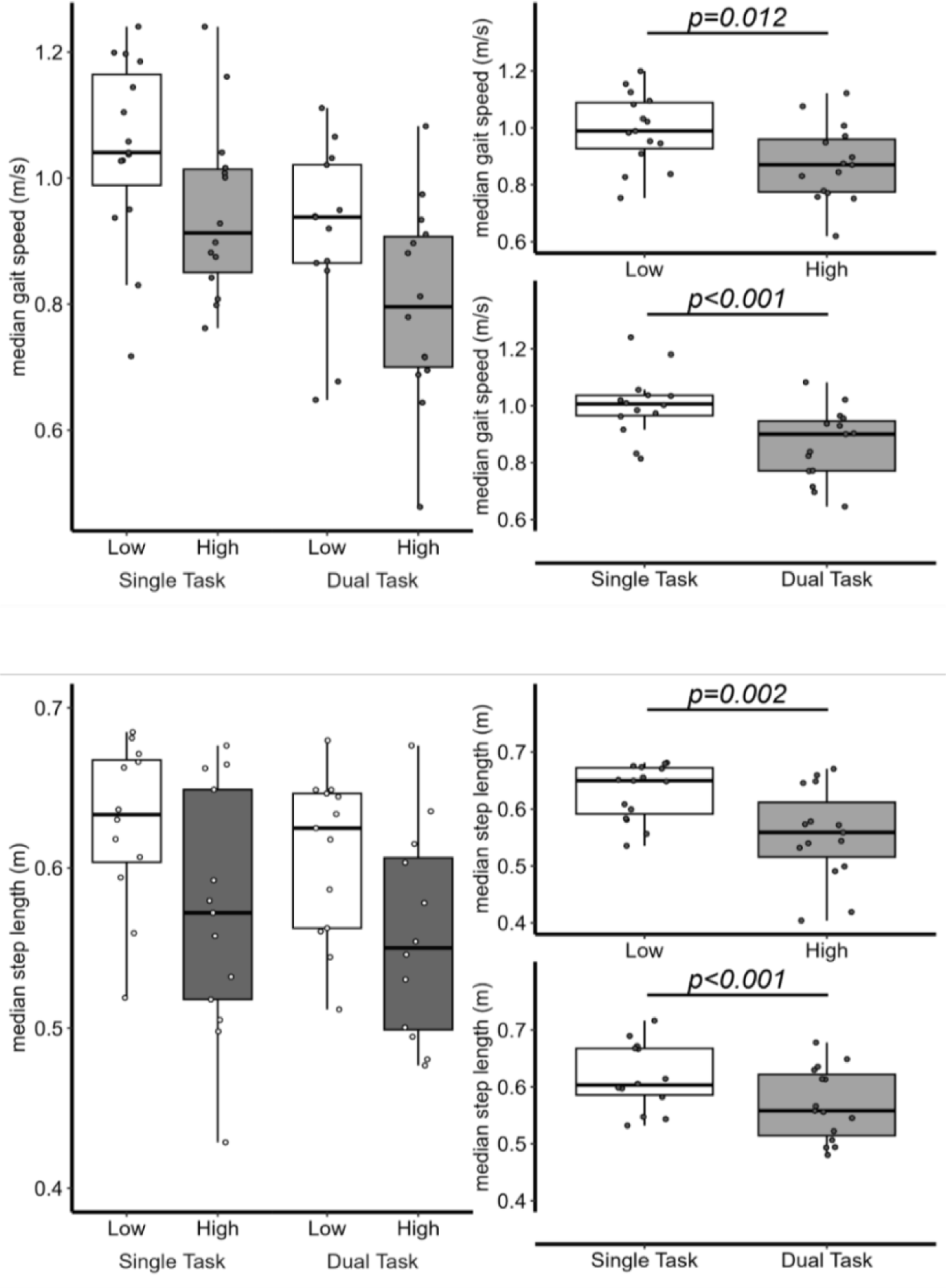
The median gait speed (top) interaction (top left) and main effects (top right) and step length (bottom) and main effects (bottom right) across virtual Low versus High elevation and Single versus Dual-task

We documented significant main effects of Height, *F*(1,52) = 10.02, *p* = 0.003, and Cognitive Demand, *F*(1,52) = 15.39, *p* < 0.001, for step length, revealing that participants shortened their steps approximately 11% during the high compared to low elevation condition (*β* = -0.07 m [-0.11,-0.03]) and 7.3% during dual-task vs. single task walking (*β* = -0.05 m [-0.07,-0.02]), but no interactions were documented for step length (*p* = 0.427) and no main effects or interactions were revealed for step length variability (all *p*’s > .179; see Figure 3 and Supplementary Tables 7-8).

Models analyzing walking condition effects on step width revealed a significant main effect of Cognitive Demand, *F*(1,52) = 6.35, *p* = 0.015, with participants adopting a more conservative (i.e., 7.7% wider) stepping pattern during dual task walking (*β* = 0.01 m [0.003,0.02]). No significant main effects of Height or Height x Cognitive Demand interactions were observed for step width (all *p*’s > .301). Finally, analyses of step width variability revealed a significant main effect of height, *F*(1,52) = 12.60, *p* < 0.001, such that participants exhibited less variable step width at high compared to low virtual elevation (*β* = -0.009 m [-0.01,-0.004]), but no other main effects or interactions were observed (all *p*’s > .264; see Figure 3 and Supplementary Tables 9-10).

**Figure 4.**
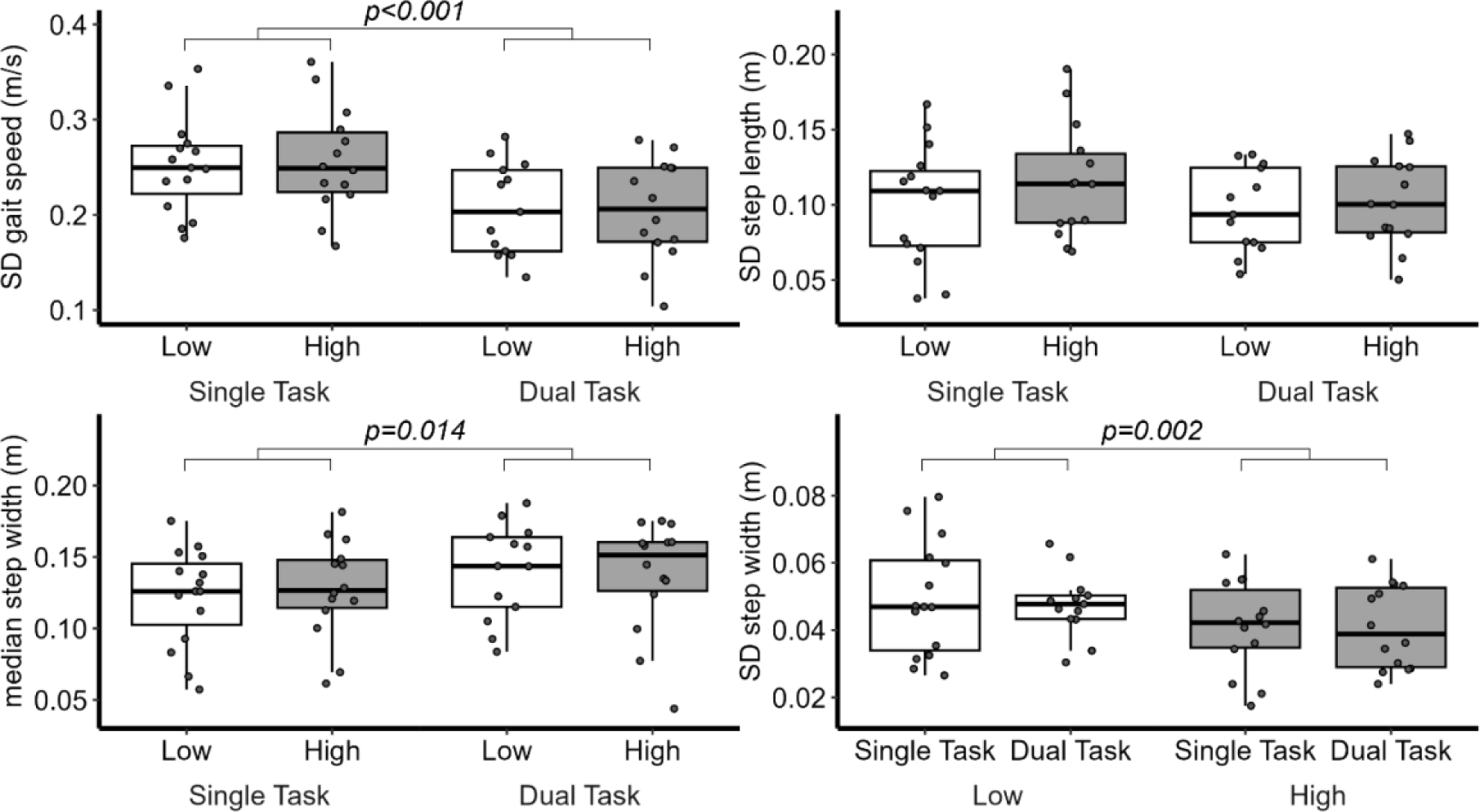
The standard deviation (SD) of gait speed (Top Left), step length (Top Right), and mean step width (Bottom Left), and SD step width (Bottom Right) for each walking condition.

### 3.3 Extemporaneous Speech Performance

For the concurrent extemporaneous speech task, a main effect of Task-Elevation was observed across all silent speech pause time in seconds, *F*(2,948) = 3.719, *p* = 0.024. Follow-up planned pairwise comparisons (with Bonferroni corrections) revealed that silent speech pauses were longer while walking in the high (+ 17.8%, *p* < 0.001) and low conditions (+ 2.8%, *p* = 0.047) compared to single-task seated performance, but pauses were not significantly different between virtual low and high elevation conditions (*p* = 0.196). There were no main effects for the number of speech pauses, *F*(2,35) = 0.341, *p* = 0.714, or for the total speech pause duration, *F*(2,35) = 1.385, *p* = 0.264.

**Figure 5.**
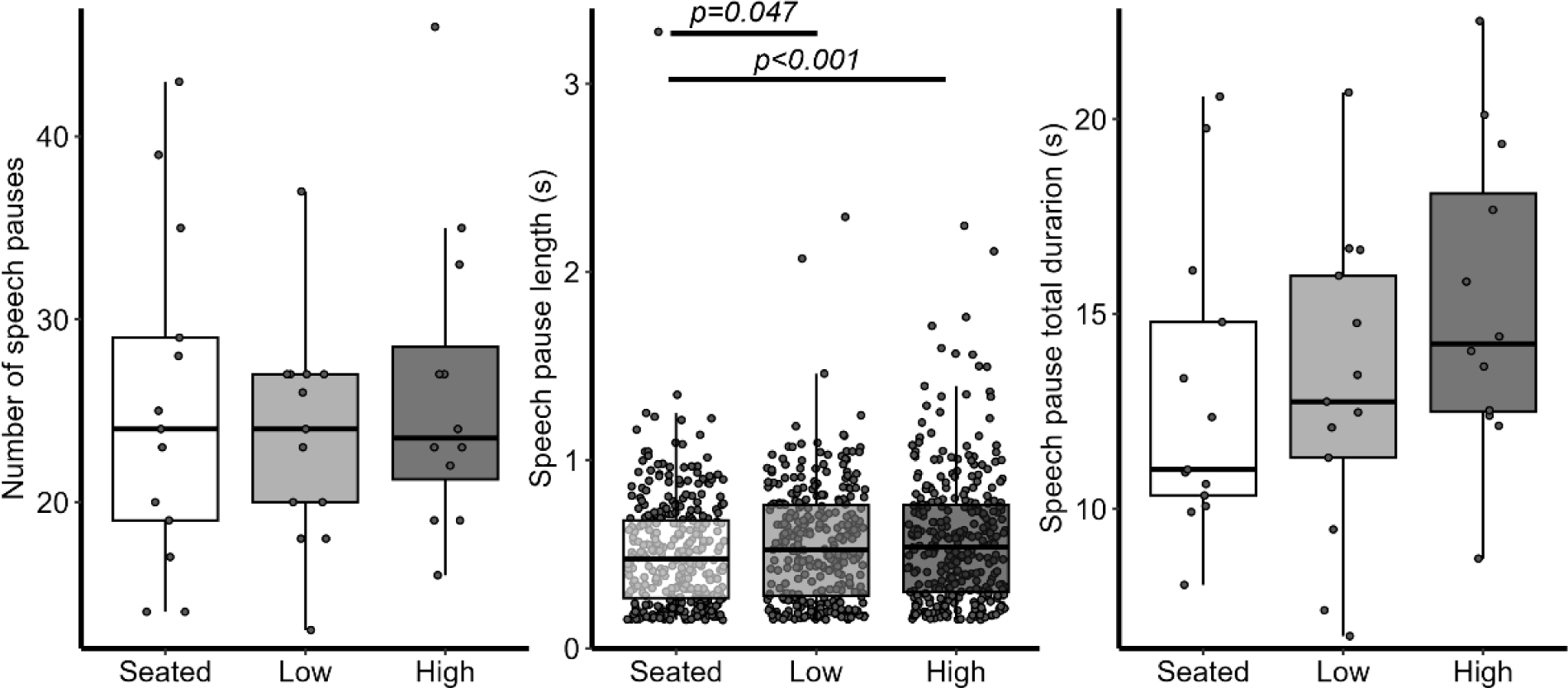
The extemporaneous speech mean silent pause number (left), length (middle), and total duration of silent pauses for each condition (right).

## 4. Discussion

The present study used a familiar, but challenging, cognitive task (extemporaneous speech) to study how healthy adults managed dual-task costs during mobility in environments of varying levels of balance threat. We expected that extemporaneous speech would lead to cognitive-motor interference that would be reflected in an interaction between cognitive demands and mobility-related anxiety while participants were walking and talking at virtual high elevation. We did not detect an interaction between cognitive demands and mobility-related anxiety as predicted, but main effects were revealed across all performance outcomes. Findings from speech pauses (i.e., measure of cognitive interference) suggest healthy young adults prioritize talking while seated or walking, even when walking behavior is threatened by a virtual elevation. A lack of interaction effects detected suggest independent biobehavioral mechanisms are at play when walking and concurrently engaging in cognitively demanding tasks that are well-practiced and rely on different sensorimotor processes (e.g., verbal, self-generated) as the motor task (e.g., visual, proprioceptive). Gait performance was adjusted similarly to cope with cognitive demand (slower speeds, shorter and wider steps) or added mobility-related anxiety (slower speeds, shorter steps), with the exception of increases in step-width that were exclusively observed during extemporaneous speech.

Results further suggest that dual-task demands may be prioritized in some situations over walking, even when walking under conditions that threaten safety. Data from the extemporaneous task indicated that participants demonstrated an increase in silent speech pauses, indicative of increased cognitive interference, from seated to walking. However, we detected no substantively greater cognitive interference as a result of state mobility-related anxiety, despite an established negative association between anxiety and speech performance.^53,54^ We interpret this finding as participants ensuring sufficient resources were available to devote to a cognitive task when it is well-practiced like extemporaneous speech and not directly interfering with the motor task, thereby allowing cognitive-motor costs to be reflected in mobility rather than their speech production. Previous work suggests young adults do not need to prioritize the gait task until motor complexity is challenged (i.e. obstacle avoidance,^4^ narrower path^27^). It is possible that the high elevation environment was not threatening enough for healthy and capable individuals to sacrifice speech in favor of their walking performance.^2^ As healthy adults, they may have been suitably confident that they would not fall in the virtual environment.^14^ We suspect the results may change if the cognitive or motor demands were greater at high elevation, such as walking at a faster speed ^55–57^ or avoiding an obstacle.^4,58^ We expect a population that is less confident in their capacity to perform both tasks adequately (e.g., older adults,^59,60^ individuals with motor impairments) ^61,62^ would demonstrate different prioritization strategies.

Self-report data from Likert scales provided within each series of trial blocks indicated that participants were more anxious, less confident, and devoted more effort towards the task when walking at elevation compared to ground level and when performing the walking task while talking compared to without talking. Compared to single-task conditions, walking while talking was not associated with significant changes in participants’ self-reported anxiety, confidence, or mental effort. Although not a significant effect, there was an interaction trend in the data observed for cognitive anxiety (i.e., worrisome thoughts, *p* = .074) and self-reported somatic anxiety (i.e. perceptions of changes in arousal, *p* = .073), suggesting that people were less perceptive of changes in their physiological reaction to the balance threat at high elevation while talking compared to without the speaking. In alignment with this interpretation, participants’ self-reported mental effort when walking at high elevation was not sensitive to the addition of a concurrent extemporaneous speech task, supporting that walking at high elevation with a cognitive task recruited no perceivably greater mental effort than only walking in anxiety-inducing settings.

In alignment with previous research imposing similar virtual balance threat during mobility tasks there was no effect on step width at high elevation,^35,60^ although a significant increase in step width and step width variability during the dual-task was evidenced. Previously, researchers have reported an increase in step width during a dual-task,^58^ which is interpreted as a means of gaining added mediolateral stability during a cognitively demanding activity.^63^ In the current paper, the primary difference between walking at high elevation and walking with a cognitive demand is that balance threat encourages a narrower stepping pattern, but the inclusion of a dual task encourages slower gait that takes the form of slower, somewhat shorter, and wider (albeit variably) steps. As a result of fixed platform dimensions in the current study, increasing step width at high elevation would bring feet closer to the edge of the walkway and increase the probability of a potential fall.^34^ Apparently, our participants coped with added cognitive demands by reducing gait speed and gait speed variability at the same time as widening their steps, but competing step width goals at virtual high elevation prevented an interaction from being revealed in gait performance.

Given the lack of interaction effects for measures of gait performance, the results suggest that instead of a conflicting resource demand, cognitive-motor resources involved in gait may tap predominantly independent processes to those required to cope with state anxiety. Therefore, the combination of cognitive-motor demands and state anxiety may not compound the deleterious effects on walking behavior. Alternatively, the act of engaging in extemporaneous speech could be a distraction from reinvestment or rumination when balance is threatened. Anecdotally, participants frequently commented that “it was actually easier to walk at high heights while talking” after study participation. The interpretation that speaking was a distraction that benefitted motor performance aligns with evidence of the positive benefits of self-talk as a coping skill, even when speech is not directed at the primary motor task.^64,65^ Cognitive-motor demands serving as a distractor would align with existing theoretical assumptions about cognitive and attentional processes under anxiety,^15,16^ as well as empirical evidence from studies using dual-task gait paradigms in healthy young ^38^ and older adults.^18^ Focusing on the cognitive task could have allowed self-organization processes to control walking without interference, allowing gait behavior to unfold implicitly and attention to be focused externally. However, given somewhat conflicting results between self-report measures and gait performance in the present data, further research should pursue the results observed here by including a more impaired population that would be less capable of coping with concurrent mobility-related anxiety and cognitive-motor demands.

### 4.1 Limitations and Future Directions

In the future, researchers should aim to further validate these findings among a larger and more diverse sample, as well as extending this paradigm to populations with movement impairments or psychological traits (e.g., trait anxiety) that could elicit greater sensitivity to cognitive and motor demands. Indirect measures of attentional allocation (e.g., through the use of gaze tracking) might better clarify how individuals extract information during ambulation while performing concurrent cognitive tasks. Similarly, subjective indices of attentional and motor resource allocation using recently developed self-report instruments might further enhance understanding of prioritization during complex cognitive and mobility tasks.^66^ Extemporaneous speech topics, while familiar and accessible to participants, may possess some inherent affective content (e.g., a pleasant memory of time spent with a friend). In future studies, researchers should explore the affective context of the speech monologue to potentially control for confounds from affect-induced changes in gait behavior.^67,68^ Finally, measuring lexical complexity and lexical ‘stageholders’ like filled pauses in extemporaneous speech should be included in future studies.^69^

### 4.2 Summary and Conclusions

In conclusion, we built on previous evidence of attentional control under anxiety by testing an ecologically relevant and well-practiced cognitive task: extemporaneous speech. We successfully induced mobility-related anxiety in healthy young adults using a virtual balance threat, evidenced by decreases in self-reported confidence and increases in anxiety and mental effort. Gait kinematics indicate that, compared to ground level, walking at simulated elevation is associated with participants adopting a more conservative gait (i.e., slower, shorter steps, with less variability in step width). Participants prioritized the extemporaneous speech task by walking slower (and with less variability) and taking shorter and wider steps. However, no interaction effects in gait behavior were documented, suggesting that the well-practiced cognitive-motor demands of talking were not additive to the effects of mobility-related state anxiety on the locomotor system. Speech pause duration and number were affected by motor complexity but were seemingly unaffected by the virtual mobility threat. Trends in participants’ subjective feelings of worry and confidence during the task, along with informal debriefing, suggest talking while walking may buffer individuals from the deleterious effects of anxiety on mobility through distraction.

## Acknowledgements

We thank our participants who volunteered their time as we were just emerging from the pandemic. We thank Dr. Jessica Huber, Caitlin Kane, and Mackenzie Barrowman, for their guidance and effort analyzing the extemporaneous speech data. We thank Benjamin Engle and the VR lab at the University of Utah Eccles Health Sciences Library for their craftwork designing and refining the VR program we used in this study. This study was made possible through the support of the Office of Undergraduate Research Opportunity Program and a Research Incentive Award at the University of Utah.

## Notes

### Competing Interest Statement

The authors have declared no competing interest.

## References

1. Woollacott M, Shumway-Cook A. Attention and the control of posture and gait: A review of an emerging area of research. Gait and Posture. 2002;16(1):1–14. doi:10.1016/S0966-6362(01)00156-4

2. Yogev-Seligmann G, Hausdorff JM, Giladi N. Do we always prioritize balance when walking? Towards an integrated model of task prioritization. Movement Disorders. 2012;27(6):765–770. doi:10.1002/mds.24963

3. Siu KC, Catena RD, Chou LS, van Donkelaar P, Woollacott MH. Effects of a secondary task on obstacle avoidance in healthy young adults. Experimental Brain Research. 2008;184(1):115–120. doi:10.1007/s00221-007-1087-9

4. Raffegeau TE, Haddad JM, Huber JE, Rietdyk S. Walking while talking: Young adults flexibly allocate resources between speech and gait. Gait & Posture. 2018;64:59–62. doi:10.1016/j.gaitpost.2018.05.029

5. Beurskens R, Bock O. Does the walking task matter? Influence of different walking conditions on dual-task performances in young and older persons. Human Movement Science. 2013;32(6):1456–1466. doi:10.1016/j.humov.2013.07.013

6. Beurskens R, Helmich I, Rein R, Bock O. Age-related changes in prefrontal activity during walking in dual-task situations: a fNIRS study. Int J Psychophysiol. 2014;92:122–128. doi:10.1016/j.ijpsycho.2014.03.005

7. Holtzer R, Mahoney JR, Izzetoglu M, Izzetoglu K, Onaral B, Verghese J. fNIRS study of walking and walking while talking in young and old individuals. Journals of Gerontology - Series A Biological Sciences and Medical Sciences. 2011;66 A(8):879-887. doi:10.1093/gerona/glr068

8. Holtzer R, Izzetoglu M. Mild cognitive impairments attenuate prefrontal cortex activations during walking in older adults. Brain Sciences. 2020;10(7):1–16. doi:10.3390/brainsci10070415

9. Montero-Odasso MM, Sarquis-Adamson Y, Speechley M, et al. Association of dual-task gait with incident dementia in mild cognitive impairment: Results from the gait and brain study. JAMA Neurology. 2017;74(7):857–865. doi:10.1001/jamaneurol.2017.0643

10. Li F, Harmer P. Prevalence of falls, physical performance, and dual-task cost while walking in older adults at high risk of falling with and without cognitive impairment. Clinical Interventions in Aging. 2020;15:945–952. doi:10.2147/CIA.S254764

11. Baker K, Rochester L, Nieuwboer A. The Immediate Effect of Attentional, Auditory, and a Combined Cue Strategy on Gait During Single and Dual Tasks in Parkinson’s Disease. Archives of Physical Medicine and Rehabilitation. 2007;88(12):1593–1600. doi:10.1016/j.apmr.2007.07.026

12. Beck EN, Intzandt BN, Almeida QJ. Can Dual Task Walking Improve in Parkinson’s Disease After External Focus of Attention Exercise? A Single Blind Randomized Controlled Trial. Neurorehabilitation and Neural Repair. 2018;32(1):18–33. doi:10.1177/1545968317746782

13. Camicioli R, Oken BS, Sexton G, Kaye JA, Nutt JG. Verbal fluency task affects gait in Parkinson’s disease with motor freezing. Journal of Geriatric Psychiatry and Neurology. 1998;11(4):181–185. doi:10.1177/089198879901100403

14. Young WR, Williams AM. How fear of falling can increase fall-risk in older adults: Applying psychological theory to practical observations. Gait and Posture. 2015;41(1):7–12. doi:10.1016/j.gaitpost.2014.09.006

15. Eysenck MW, Derakshan N, Santos R, Calvo MG. Anxiety and cognitive performance: Attentional Control Theory. Emotion. 2007;7(2):336–353. doi:10.1037/1528-3542.7.2.336

16. Masters R, Maxwell J. The theory of reinvestment. International Review of Sport and Exercise Psychology. 2008;1(2):160–183. doi:10.1080/17509840802287218

17. Uiga L, Capio CM, Ryu D, et al. The Role of Movement-Specific Reinvestment in Visuomotor Control of Walking by Older Adults. The Journals of Gerontology: Series B. Published online June 2018:1–11. doi:10.1093/geronb/gby078

18. Young WR, Olonilua M, Masters RSW, Dimitriadis S, Williams AM. Examining links between anxiety, reinvestment and walking when talking by older adults during adaptive gait. Experimental Brain Research. 2016;234(1):161–172. doi:10.1007/s00221-015-4445-z

19. Ellmers TJ, Cocks AJ, Kal EC, Young WR. Conscious Movement Processing, Fall-Related Anxiety, and the Visuomotor Control of Locomotion in Older Adults. Taler V, ed. The Journals of Gerontology: Series B. 2020;75(9):1911–1920. doi:10.1093/geronb/gbaa081

20. Wulf G. Attentional focus and motor learning: a review of 15 years. International Review of Sport and Exercise Psychology. 2013;6(1):77–104. doi:10.1080/1750984X.2012.723728

21. Kal EC, Young WR, Ellmers TJ. Balance capacity influences the effects of conscious movement processing on postural control in older adults. Human Movement Science. 2022;82(January):102933. doi:10.1016/j.humov.2022.102933

22. Nnodim JO, Kim H, Ashton-Miller JA. Dual-task performance in older adults during discrete gait perturbation. Experimental Brain Research. 2016;234(4):1077–1084. doi:10.1007/s00221-015-4533-0

23. Plummer-D’Amato P, Altmann LJP, Reilly K. Dual-task effects of spontaneous speech and executive function on gait in aging: Exaggerated effects in slow walkers. Gait and Posture. 2011;33(2):233–237. doi:10.1016/j.gaitpost.2010.11.011

24. Worden TA, Vallis LA. Concurrent performance of a cognitive and dynamic obstacle avoidance task: Influence of dual-task training. Journal of Motor Behavior. 2014;46(5):357–368. doi:10.1080/00222895.2014.914887

25. Kimura N, van Deursen R. The Effect Of Visual Dual-Tasking Interference On Walking In Healthy Young Adults. Gait & Posture. Published online April 2020. doi:10.1016/j.gaitpost.2020.04.018

26. Schaefer S, Schellenbach M, Lindenberger U, Woollacott MH. Walking in high-risk settings: do older adults still prioritize gait when distracted by a cognitive task? Exp Brain Res. 2015;233(1):79–88. doi:10.1007/s00221-014-4093-8

27. Lindenberger U, Marsiske M, Baltes PB. Memorizing while walking: Increase in dual-task costs from young adulthood to old age. Psychology and Aging. 2000;15(3):417–436. doi:10.1037/0882-7974.15.3.417

28. Birnie K, Martin RM, Gallacher J, et al. Socio-economic disadvantage from childhood to adulthood and locomotor function in old age: a lifecourse analysis of the Boyd Orr and Caerphilly prospective studies. J Epidemiol Community Health. 2011;65(11):1014–1023. doi:10.1136/jech.2009.103648

29. Penati R, Schieppati M, Nardone A. Cognitive performance during gait is worsened by overground but enhanced by treadmill walking. Gait & Posture. 2020;76:182–187. doi:10.1016/j.gaitpost.2019.12.006

30. Yogev G, Giladi N, Peretz C, Springer S, Simon ES, Hausdorff JM. Dual tasking, gait rhythmicity, and Parkinson’s disease: Which aspects of gait are attention demanding? European Journal of Neuroscience. 2005;22(5):1248–1256. doi:10.1111/j.1460-9568.2005.04298.x

31. Miyake A, Friedman NP, Emerson MJ, Witzki AH, Howerter A. The unity and diversity of executive functions and their contributions to complex “frontal lobe” tasks: a latent variable analysis. Cogn Psychol. 2000;41:49–100. doi:10.1006/cogp.1999.0734

32. Lovett MC. A Strategy-Based Interpretation of Stroop. Cognitive Science. 2005;29(3):493–524. doi:10.1207/s15516709cog0000_24

33. Raffegeau TE, Fawver B, Clark M, et al. The feasibility of using virtual reality to induce mobility-related anxiety during turning. Gait & Posture. 2020;77:6–13. doi:10.1016/j.gaitpost.2020.01.006

34. Raffegeau TE, Fawver B, Young WR, Williams AM, Lohse KR, Fino PC. The direction of postural threat alters balance control when standing at virtual elevation. Experimental Brain Research. 2020;238(11):2653–2663. doi:10.1007/s00221-020-05917-5

35. Raffegeau TE, Clark M, Fawver B, et al. The effect of mobility-related anxiety on walking across the lifespan: A virtual reality simulation study. Experimental Brain Research. 2023;19:1–12. doi:doi: 10.1007/s00221-023-06638-1

36. Cleworth TW, Horslen BC, Carpenter MG. Influence of real and virtual heights on standing balance. Gait & Posture. 2012;36(2):172–176. doi:10.1016/j.gaitpost.2012.02.010

37. Adkin AL, Carpenter MG. New insights on emotional contributions to human postural control. Frontiers in Neurology. 2018;9. doi:10.3389/fneur.2018.00789

38. Ellmers TJ, Young WR. Conscious motor control impairs attentional processing efficiency during precision stepping. Gait and Posture. 2018;63(April):58–62. doi:10.1016/j.gaitpost.2018.04.033

39. Ellmers TJ, Cocks AJ, Kal EC, Young WR. Conscious movement processing, fall-related anxiety, and thevisuomotor control of locomotion in older adults. Journals of Gerontology - Series B Psychological Sciences and Social Sciences. 2020;75(9):1911–1920. doi:10.1093/geronb/gbaa081

40. Eysenck MW, Calvo MG. Anxiety and performance: the Processing Efficiency Theory. Cognition & Emotion. 1992;6(6):409–434. doi:10.1080/02699939208409696

41. Montero-Odasso M, van der Velde N, Martin FC, et al. World guidelines for falls prevention and management for older adults: A global initiative. Age and Ageing. 2022;51(9):afac205. doi:10.1093/ageing/afac205

42. Spielberger CD. State-Trait Anxiety Inventory for Adults (STAI-AD). Published online 1983. 10.1037/t06496-000

43. Raffegeau TE, Kellaher GK, Terza MJ, Roper JA, Altmann LJ, Hass CJ. Older women take shorter steps during backwards walking and obstacle crossing. Experimental Gerontology. 2019;122:60–66. doi:10.1016/j.exger.2019.04.011

44. Hobert MA, Niebler R, Meyer SI, et al. Poor Trail Making Test Performance Is Directly Associated with Altered Dual Task Prioritization in the Elderly – Baseline Results from the TREND Study. Laks J, ed. PLoS ONE. 2011;6(11):e27831. doi:10.1371/journal.pone.0027831

45. Arbuthnott K, Frank J. Trail Making Test, Part B as a Measure of Executive Control : Validation Using a Set-Switching Paradigm Trail Making Test, Part B as a Measure of Executive Control : Validation Using a Set-Switching Paradigm *. 2010;3395(915545541). doi:10.1076/1380-3395(200008)22

46. Jensen AR, Rohwer WD. The Stroop Color-Word test: a review. Acta Psychologica. 1966;25(C):36–93. doi:10.1016/0001-6918(66)90004-7

47. Krane V. The Mental Readiness Form as a Measure of Competitive State Anxiety. The Sport Psychologist. 2016;8(2):189–202. doi:10.1123/tsp.8.2.189

48. Zijlstra FRH. Efficiency in Work Behaviour: A Design Approach for Modern Tools. Delft University Press; 1993.

49. International Society of Biomechanics Standardization and Terminology Committee. ISB recommendation on definitions of joint coordinate system of various joints for the reporting of human joint motion-part I: ankle, hip, and spine. Journal of Biomechanics. 2002;(35):543-548. doi:10.1300/J181v01n04_07

50. Zeni J, Richards J, Higginson JS. Two simple methods for determining gait events during treadmill and overground walking using kinematic data. Gait Posture. 2008;27(4):710–714. doi:10.1016/j.gaitpost.2007.07.007

51. Darling-White M, Huber JE. The Impact of Parkinson’s Disease on Breath Pauses and Their Relationship to Speech Impairment: A Longitudinal Study. American Journal of Speech-Language Pathology. Published online July 2020:1–13. doi:10.1044/2020_AJSLP-20-00003

52. Lee J, Huber J, Jenkins J, Fredrick J. Language planning and pauses in story retell: Evidence from aging and Parkinson’s disease. Journal of Communication Disorders. 2019;79:1–10. doi:10.1016/J.JCOMDIS.2019.02.004

53. Lay CH, Paivio A. The effects of task difficulty and anxiety on hesitations in speech. Canadian Journal of Behavioural Science / Revue canadienne des sciences du comportement. 1969;1(1):25–37. doi:10.1037/h0082683

54. Laukka P, Linnman C, Åhs F, et al. In a Nervous Voice: Acoustic Analysis and Perception of Anxiety in Social Phobics’ Speech. J Nonverbal Behav. 2008;32(4):195–214. doi:10.1007/s10919-008-0055-9

55. Callisaya ML, Launay CP, Srikanth VK, Verghese J, Allali G, Beauchet O. Cognitive status, fast walking speed and walking speed reserve—the Gait and Alzheimer Interactions Tracking (GAIT) study. GeroScience. 2017;39(2):231–239. doi:10.1007/s11357-017-9973-y

56. Dennis A, Dawes H, Elsworth C, et al. Fast walking under cognitive-motor interference conditions in chronic stroke. Brain Research. 2009;1287:104–110. doi:10.1016/j.brainres.2009.06.023

57. Patel P, Lamar M, Bhatt T. Effect of type of cognitive task and walking speed on cognitive-motor interference during dual-task walking. Neuroscience. 2014;260:140–148. doi:10.1016/j.neuroscience.2013.12.016

58. Raffegeau TE, Brinkerhoff SA, Kellaher GK, et al. Changes to margins of stability from walking to obstacle crossing in older adults while walking fast and with a dual-task. Experimental Gerontology. 2022;161(October 2021):111710. doi:10.1016/j.exger.2022.111710

59. Brown LA, Gage WH, Polych MA, Sleik RJ, Winder TR. Central set influences on gait: Age-dependent effects of postural threat. Experimental Brain Research. 2002;145(3):286–296. doi:10.1007/s00221-002-1082-0

60. Gage WH, Sleik RJ, Polych MA, McKenzie NC, Brown LA. The allocation of attention during locomotion is altered by anxiety. Exp Brain Res. 2003;150:385–394. doi:10.1007/s00221-003-1468-7

61. Ehgoetz Martens KA, Silveira CRA, Intzandt BN, Almeida QJ. Overload From Anxiety: A Non-Motor Cause for Gait Impairments in Parkinson’s Disease. The Journal of Neuropsychiatry and Clinical Neurosciences. Published online 2017:appi.neuropsych. doi:10.1176/appi.neuropsych.16110298

62. Ehgoetz-Martens KA, Ellard CG, Almeida QJ. Virtually-induced threat in Parkinson’s: Dopaminergic interactions between anxiety and sensory–perceptual processing while walking. Neuropsychologia. 2015;79:322–331. doi:10.1016/j.neuropsychologia.2015.05.015

63. Bauby CE, Kuo AD. Active control of lateral balance in human walking. J Biomech. 2000;33:1433–1440.

64. Walter N, Nikoleizig L, Alfermann D. Effects of Self-Talk Training on Competitive Anxiety, Self-Efficacy, Volitional Skills, and Performance: An Intervention Study with Junior Sub-Elite Athletes. Sports. 2019;7(6):148. doi:10.3390/sports7060148

65. Hatzigeorgiadis A, Zourbanos N, Mpoumpaki S, Theodorakis Y. Mechanisms underlying the self-talk–performance relationship: The effects of motivational self-talk on self-confidence and anxiety. Psychology of Sport and Exercise. 2009;10(1):186–192. doi:10.1016/j.psychsport.2008.07.009

66. Young WR, Ellmers TJ, Kinrade N, Cossar J, Cocks AJ. Re-evaluating the measurement and influence of conscious movement processing on gait performance in older adults: development of the Gait-Specific Attentional Profile. Gait & Posture. 2020;81:73–77. doi:10.1016/j.gaitpost.2020.07.008

67. Fawver B, Hass CJ, Park KD, Janelle CM. Autobiographically recalled emotional states impact forward gait initiation as a function of motivational direction. Emotion. 2014;14(6):1125–1136. doi:10.1037/a0037597

68. Fawver B, Hass CJ, Coombes SA, Trapp SK, Janelle CM. Recalling fearful memories modifies approach and avoidance behavior based on spatial context. Emotion. 2022;22(3):430–443. doi:10.1037/emo0000940

69. Davie KL, Oram Cardy JE, Holmes JD, et al. The effects of word length, articulation, oral-motor movement, and lexicality on gait: A pilot study. Gait and Posture. 2012;35(4):691–693. doi:10.1016/j.gaitpost.2011.12.006

