## Supplemental Data Table 1-10 for "Walking (and talking) the plank: Dual-task performance costs in a virtual balance-threatening environment"

### Supplementary Data Tables

**Supplemental Table 1. LMER model regression: Cognitive anxiety**

| Fixed effects | *DoF* | *F* | *p* | $\beta$ | Lower limit | Upper limit |
|---|---|---|---|---|---|---|
| **Intercept** | **1, 56** | **19.09** | **<0.001** | **1.93** | **1.02** | **2.83** |
| ***Height*** | **1, 56** | **64.32** | **<0.001** | **3.42** | **2.53** | **4.32** |
| *Cognitive Demand* | 1, 56 | 2.46 | 0.267 | 0.68 | -0.33 | 1.69 |
| *Height × Cognitive Demand* | 1, 56 | 11.00 | 0.074 | -0.89 | -0.33 | -0.29 |
| Random effects | | | | $\beta$ | Lower limit | Upper limit |
| *Subject (std)* | | | | 0.003 | <0.001 | 0.007 |
| *Height:Subject (std)* | | | | 0.67 | <0.001 | 1.34 |
| *Cognitive Demand:Subject (std)* | | | | 0.85 | <0.001 | 1.70 |
| Model Error *(std)* | | | | 0.45 | 0.34 | 0.57 |

*Note:* Significant tests are bolded; *DoF* = degrees of freedom; *F* = Type 2 ANOVA test *F* value; *p* = probability value for significance test; $\beta$ = unstandardized beta weight, Lower limit and Upper limit = 95% confidence interval, std = standard deviation.

**Supplemental Table 2. LMER model regression: Somatic anxiety**

| Fixed effects | *DoF* | *F* | *p* | $\beta$ | Lower limit | Upper limit |
|---|---|---|---|---|---|---|
| **Intercept** | **1, 56** | **32.65** | **<0.001** | **2.73** | **1.75** | **3.71** |
| ***Height*** | **1, 56** | **85.03** | **<0.001** | **3.28** | **2.55** | **4.02** |
| *Cognitive Demand* | 1, 56 | 1.00 | 0.215 | -0.65 | -0.65 | 1.08 |
| *Height × Cognitive Demand* | 1, 56 | 8.11 | 0.190 | -0.76 | -0.65 | -0.02 |
| Random effects | | | | $\beta$ | Lower limit | Upper limit |
| *Subject (std)* | | | | 0.88 | <0.001 | 1.76 |
| *Height:Subject (std)* | | | | 0.32 | <0.001 | 0.63 |
| *Cognitive Demand:Subject (std)* | | | | 0.51 | <0.001 | 1.02 |
| Model Error *(std)* | | | | 0.59 | 0.43 | 0.74 |

*Note:* Significant tests are bolded; *DoF* = degrees of freedom; *F* = Type 2 ANOVA test *F* value; *p* = probability value for significance test; $\beta$ = unstandardized beta weight, Lower limit and Upper limit = 95% confidence interval, std = standard deviation.

**Supplemental Table 3. LMER model regression: Confidence**

| Fixed effects | *DoF* | *F* | *p* | *β* | Lower limit | Upper limit |
|---|---|---|---|---|---|---|
| **Intercept** | **1, 56** | **578.82** | **<0.001** | **10.19** | **9.28** | **11.11** |
| ***Height*** | **1, 56** | **53.81** | **<0.001** | **-2.88** | **-3.77** | **-1.98** |
| *Cognitive Demand* | 1, 56 | 1.76 | 0.261 | -0.60 | -1.56 | 0.36 |
| *Height × Cognitive Demand* | 1, 56 | 18.02 | 0.073 | 0.94 | -1.56 | 1.52 |

| Random effects | *β* | Lower limit | Upper limit |
|---|---|---|---|
| *Subject (std)* | 0.13 | <.001 | 0.26 |
| *Height:Subject (std)* | 0.69 | <.001 | 1.38 |
| *Cognitive Demand:Subject (std)* | 0.79 | <.001 | 1.59 |
| Model Error *(std)* | 0.42 | 0.31 | 0.53 |

*Note:* Significant tests are bolded; *DoF* = degrees of freedom; *F* = Type 2 ANOVA test *F* value; *p* = probability value for significance test; *β* = unstandardized beta weight, Lower limit and Upper limit = 95% confidence interval, std = standard deviation.

**Supplemental Table 4. LMER model regression: Mental Effort**

| Fixed effects | *DoF* | *F* | *p* | *β* | Lower limit | Upper limit |
|---|---|---|---|---|---|---|
| **Intercept** | **1, 56** | **16.69** | **0.003** | **20.80** | **9.34** | **32.27** |
| ***Height*** | **1, 56** | **35.60** | **<0.001** | **29.40** | **16.07** | **42.72** |
| *Cognitive Demand* | 1, 56 | 4.75 | 0.111 | 8.00 | 0.05 | 15.95 |
| *Height × Cognitive Demand* | 1, 56 | 3.84 | 0.353 | -0.77 | 0.05 | 5.79 |

| Random effects | *β* | Lower limit | Upper limit |
|---|---|---|---|
| *Subject (std)* | 3.17 | <0.001 | 6.35 |
| *Height:Subject (std)* | 12.09 | <0.001 | 24.18 |
| *Cognitive Demand:Subject (std)* | 4.72 | <0.001 | 9.45 |
| Model Error *(std)* | 4.91 | 3.64 | 6.17 |

*Note:* Significant tests are bolded; *DoF* = degrees of freedom; *F* = Type 2 ANOVA test *F* value; *p* = probability value for significance test; *β* = unstandardized beta weight, Lower limit and Upper limit = 95% confidence interval, std = standard deviation.

**Supplemental Table 5. LMER model regression: Gait speed (m/s)**

| Fixed effects | *DoF* | *F* | *p* | *β* | Lower limit | Upper limit |
|---|---|---|---|---|---|---|
| **Intercept** | **1, 52** | **863.37** | **< 0.001** | **1.07** | **0.99** | **1.14** |
| ***Height*** | **1, 52** | **7.12** | **0.010** | **-0.11** | **-0.20** | **-0.03** |
| ***Cognitive Demand*** | **1, 52** | **32.11** | **< 0.001** | **-0.16** | **-0.21** | **-0.10** |
| *Height × Cognitive Demand* | 1, 52 | 0.004 | 0.951 | 0.002 | -0.06 | 0.07 |
| Random effects | | | | *β* | Lower limit | Upper limit |
| *Subject (std)* | | | | 0.066 | 0.023 | 0.190 |
| *Subject:Height (std)* | | | | 0.100 | 0.066 | 0.154 |
| *Subject:Cognitive Demand (std)* | | | | 0.038 | 0.015 | 0.095 |
| Model Error *(std)* | | | | 0.058 | 0.040 | 0.084 |

*Note:* Significant tests are bolded; m/s = meters/second, *DoF* = degrees of freedom; *F* = Type 2 ANOVA test *F* value; *p* = probability value for significance test; *β* = unstandardized beta weight, Lower limit and Upper limit = 95% confidence interval, std = standard deviation.

**Supplemental Table 6. LMER model regression: Gait speed variability (SD)**

| Fixed effects | *DoF* | *F* | *p* | *β* | Lower limit | Upper limit |
|---|---|---|---|---|---|---|
| **Intercept** | **1, 52** | **377.98** | **<0.001** | **0.27** | **0.23** | **0.28** |
| *Height* | 1, 52 | 0.06 | 0.814 | 0.004 | -0.03 | 0.04 |
| ***Cognitive Demand*** | **1, 52** | **18.21** | **<0.001** | **-0.05** | **-0.07** | **-0.03** |
| *Height × Cognitive Demand* | 1, 52 | 0.15 | 0.696 | -0.006 | -0.04 | 0.03 |
| Random effects | | | | *β* | Lower limit | Upper limit |
| *Subject (std)* | | | | 0.025 | 0.010 | 0.063 |
| *Subject:Height (std)* | | | | 0.033 | 0.019 | 0.055 |
| *Subject:Cognitive Demand (std)* | | | | <0.001 | <0.001 | <0.001 |
| Model Error *(std)* | | | | 0.030 | 0.023 | 0.039 |

*Note:* Significant tests are bolded; SD = standard deviation, *DoF* = degrees of freedom; *F* = Type 2 ANOVA test *F* value; *p* = probability value for significance test; *β* = unstandardized beta weight, Lower limit and Upper limit = 95% confidence interval, std = standard deviation.

**Supplemental Table 7. LMER model regression: Step length (m)**

| Fixed effects | *DoF* | *F* | *p* | *β* | Lower limit | Upper limit |
|---|---|---|---|---|---|---|
| **Intercept** | **1, 52** | **1293.80** | **<0.001** | **0.65** | **0.62** | **0.69** |
| ***Height*** | **1, 52** | **10.02** | **0.003** | **-0.07** | **-0.11** | **-0.03** |
| ***Cognitive Demand*** | **1, 52** | **15.39** | **<0.001** | **-0.05** | **-0.07** | **-0.02** |
| *Height × Cognitive Demand* | 1, 52 | 0. 640 | 0.427 | -0.01 | -0.04 | 0.02 |

| Random effects | *β* | Lower limit | Upper limit |
|---|---|---|---|
| *Subject (std)* | 0.034 | 0.012 | 0.094 |
| *Height:Subject (std)* | 0.053 | 0.036 | 0.079 |
| *Cognitive Demand:Subject (std)* | 0.019 | 0.009 | 0.040 |
| Model Error *(std)* | 0.024 | 0.016 | 0.034 |

*Note:* Significant tests are bolded; *m*=meters, *DoF* = degrees of freedom; *F* = Type 2 ANOVA test *F* value; *p* = probability value for significance test; *β* = unstandardized beta weight, Lower limit and Upper limit = 95% confidence interval, std = standard deviation.

**Supplemental Table 8. LMER model regression: Step length variability (SD)**

| Fixed effects | *DoF* | *F* | *p* | *β* | Lower limit | Upper limit |
|---|---|---|---|---|---|---|
| **Intercept** | **1, 52** | **139.25** | **<0.001** | **0.10** | **0.08** | **0.12** |
| *Height* | 1, 52 | 1.86 | 0.179 | 0.01 | -0.03 | 0.01 |
| *Cognitive Demand* | 1, 52 | 0.39 | 0.535 | -0.006 | -0.007 | 0.04 |
| *Height × Cognitive Demand* | 1, 52 | 0.36 | 0.549 | 0.01 | -0.04 | 0.02 |

| Random effects | *β* | Lower limit | Upper limit |
|---|---|---|---|
| *Subject (std)* | 0.015 | 0.006 | 0.037 |
| *Height:Subject (std)* | 0.014 | 0.005 | 0.039 |
| *Cognitive Demand:Subject (std)* | <0.001 | <0.001 | <0.001 |
| Model Error *(std)* | 0.025 | 0.019 | 0.033 |

*Note:* Significant tests are bolded; SD = standard deviation, *DoF* = degrees of freedom; *F* = Type 2 ANOVA test *F* value; *p* = probability value for significance test; *β* = unstandardized beta weight, Lower limit and Upper limit = 95% confidence interval, std = standard deviation.

**Supplemental Table 9. LMER model regression: Step width (m)**

| Fixed effects | *DoF* | *F* | *p* | *β* | Lower limit | Upper limit |
|---|---|---|---|---|---|---|
| **Intercept** | **1, 52** | **203.73** | **<0.001** | **0.12** | **0.11** | **0.14** |
| *Height* | 1, 52 | 0.65 | 0.424 | 0.003 | -0.006 | 0.01 |
| ***Cognitive Demand*** | **1, 52** | **6.35** | **0.015** | **0.01** | 0.003 | 0.02 |
| *Height × Cognitive Demand* | 1, 52 | 1.09 | 0.301 | -0.007 | -0.02 | 0.006 |
| Random effects | | | | *β* | Lower limit | Upper limit |
| *Subject (std)* | | | | 0.031 | 0.021 | 0.044 |
| *Height:Subject (std)* | | | | <0.001 | <0.001 | <0.001 |
| *Cognitive Demand:Subject (std)* | | | | 0.006 | 0.005 | 0.007 |
| Model Error *(std)* | | | | 0.012 | 0.010 | 0.015 |

*Note:* Significant tests are bolded; m = meters, *DoF* = degrees of freedom; *F* = Type 2 ANOVA test *F* value; *p* = probability value for significance test; *β* = unstandardized beta weight, Lower limit and Upper limit = 95% confidence interval, std = standard deviation.

**Supplemental Table 10. LMER model regression: Step width variability (SD)**

| Fixed effects | *DoF* | *F* | *p* | *β* | Lower limit | Upper limit |
|---|---|---|---|---|---|---|
| **Intercept** | **1, 52** | **223.14** | **<0.001** | **0.05** | **0.04** | **0.06** |
| ***Height*** | **1, 52** | **12.60** | **<0.001** | **-0.009** | **-0.01** | **-0.004** |
| *Cognitive Demand* | 1, 52 | 1.27 | 0.264 | -0.004 | -0.01 | 0.003 |
| *Height × Cognitive Demand* | 1, 52 | 0.78 | 0.380 | 0.003 | -0.004 | 0.01 |
| Random effects | | | | *β* | Lower limit | Upper limit |
| *Subject (std)* | | | | 0.009 | 0.006 | 0.016 |
| *Height:Subject (std)* | | | | 0.004 | 0.002 | 0.009 |
| *Cognitive Demand:Subject (std)* | | | | 0.006 | 0.003 | 0.010 |
| Model Error *(std)* | | | | 0.006 | 0.004 | 0.009 |

*Note:* Significant tests are bolded; SD = standard deviation, *DoF* = degrees of freedom; *F* = Type 2 ANOVA test *F* value; *p* = probability value for significance test; *β* = unstandardized beta weight, Lower limit and Upper limit = 95% confidence interval, std = standard deviation.
